# MTH1 deficiency selectively increases non-cytotoxic oxidative DNA damage in lung cancer cells: more bad news than good?

**DOI:** 10.1101/195750

**Authors:** Hussein H.K. Abbas, Kheloud M.H. Alhamoudi, Mark D. Evans, George D.D. Jones, Steven S. Foster

**Affiliations:** Department of Genetics and Genome Biology, University of Leicester, Leicester, Leicestershire, LE1 7RH, UK; Department of Pathology and Forensic Medicine, College of Medicine, Al-Mustansiriya University, Baghdad, Iraq; Faculty of Health and Life Sciences, De Montfort University, Leicester, Leicestershire, LE1 9BH, England

**Keywords:** Lung cancer, MTH1, NUDT1, targeted therapy, nucleotides, oxidative DNA damage, genomic instability, combined therapy, gemcitabine, cisplatin

## Abstract

**Background:** Targeted therapies are based on exploiting cancer-cell-specific genetic features or phenotypic traits to selectively kill cancer cells while leaving normal cells unaffected. Oxidative stress is a cancer hallmark phenotype. Given that free nucleotide pools are particularly vulnerable to oxidation, the nucleotide pool sanitising enzyme, MTH1, is potentially conditionally essential in cancer cells. However, findings from previous MTH1 studies have been contradictory, meaning the relevance of MTH1 in cancer is still to be determined. Here we ascertained the role of MTH1 specifically in lung cancer cell maintenance, and the potential of MTH1 inhibition as a targeted therapy strategy to improve lung cancer treatments.

**Methods:** Using siRNA-mediated knockdown or small-molecule inhibition, we tested the genotoxic and cytotoxic effects of MTH1 deficiency on H23 (p53-mutated), H522 (p53-mutated) and A549 (wildtype p53) non-small cell lung cancer cell lines relative to normal MRC-5 lung fibroblasts. We also assessed if MTH1 inhibition augments current therapies.

**Results:** MTH1 knockdown increased levels of oxidatively damaged DNA and DNA damage signaling alterations in all lung cancer cell lines but not normal fibroblasts, despite no detectable differences in reactive oxygen species levels between any cell lines. Furthermore, MTH1 knockdown reduced H23 cell proliferation. However, unexpectedly, it did not induce apoptosis in any cell line or enhance the effects of gemcitabine, cisplatin or radiation in combination treatments. Contrastingly, TH287 and TH588 MTH1 inhibitors induced apoptosis in H23 and H522 cells, but only increased oxidative DNA damage levels in H23, indicating that they kill cells independently of DNA oxidation and seemingly via MTH1-distinct mechanisms.

**Conclusions:** MTH1 has a NSCLC-specific p53-independent role for suppressing DNA oxidation and genomic instability, though surprisingly the basis of this may not be reactive-oxygen-species-associated oxidative stress. Despite this, overall our cell viability data indicates that targeting MTH1 will likely not be an across-the-board effective NSCLC therapeutic strategy; rather it induces non-cytotoxic DNA damage that could promote cancer heterogeneity and evolution.

## Introduction

Cancer cells harbour genetic mutations and exhibit phenotypic traits that are not found in normal cells. These cancer-specific features may provide avenues for the development of targeted therapies that selectively kill cancer cells [1]. The development of such strategies is a major focus of current cancer research. One such approach is synthetic lethality, which is when an acquired defect in a particular factor or pathway renders a cancer cell sensitive to inhibition of another second specific factor. Still, this approach is often limited to a certain context or cancer type, for example, inhibiting poly(ADP-ribose) polymerase (PARP) in BRCA1- and BRCA2-deficient breast and ovarian cancers [2, 3]. An alternative though related strategy that may be more effective in treating a wider range of cancers is cancer phenotypic lethality [1], which is to target factors and cell processes that are non-essential in normal cells but that become essential for cell growth following the acquisition of hallmark cancer traits [1, 4-6]. Unfortunately, until such pathways are identified and validated for therapeutic potential, radiotherapy- and chemotherapy-based treatments that are often associated with side-effects and resistance will remain the mainstay of treatments.

Oxidative stress, which arises when there is an imbalance between the production of reactive oxygen species (ROS) and the ability of a cell to counteract their levels or effects, is a hallmark cancer trait [4] that can drive both carcinogenesis and continuing tumour evolution [4, 7]. Paradoxically, ROS are the basis of much of the cytotoxicity of radiotherapy and several chemotherapy treatments. Hence, there may be several normally non-essential oxidative stress response factors and pathways that become’conditionally essential’ in cancer cells and/or significantly affect therapy responses.

ROS can react with all components of DNA to cause numerous types of lesions [8]. However, the free deoxyribonucleoside triphosphate (dNTP) pool is reportedly 190-13,000 times more susceptible to modification than DNA [9]. This suggests that a significant proportion of oxidative-stress-induced DNA damage arises via misincorporation of oxidised dNTPs during DNA replication rather than direct DNA modification. Oxidised DNA bases do not majorly disrupt DNA structure; however, they can subsequently lead to secondary types of DNA damage such as DNA single-strand breaks (SSBs) that arise when DNA glycosylases remove damaged bases during base excision repair (BER), mis-pairing events [10], and DNA double-strand breaks (DSBs) [11] through poorly defined mechanisms that could due to DNA replication stress. DSBs in particular are highly genotoxic and cytotoxic if not repaired correctly. This leads to the prediction that the pathways involved in preventing oxidised DNA base misincorporation could be critical in either promoting or suppressing cancer development and evolution depending on context [12].

Mut T Homologue 1 (MTH1) is a Nudix hydrolase family enzyme member that hydrolyses selected oxidised dNTP and NTP substrates to the corresponding mono-phosphate products and inorganic pyrophosphate to prevent their misincorporation into DNA and RNA respectively [13-15]. Primary substrates of MTH1 are dNTPs containing 8-oxo-7,8-dihydroguanine (8-oxoGua), one of the most common types of ROS-induced lesions [16], and 2-hydroxy-adenine. Supporting the idea of a role in cancer cell maintenance, MTH1 levels are elevated in various cancers [17-21], while lower MTH1 levels in U20S osteosarcoma cells and non-small cell lung cancer (NSCLC) patient samples correlates with increased levels of DNA oxidation [22, 23]. MTH1 overexpression in oncogene-expressing human cells promoted transformation [11, 24, 25], while knockdown lead to DNA-replication-associated DNA damage response (DDR) activation and senescence [11, 26]. Accordingly, the first developed MTH1 inhibitors appear to selectively inhibit cancer cell growth [22, 27]. Collectively, these findings suggest that as cells undergo malignant transformation and acquire the trait of oxidative stress, MTH1 becomes essential for maintaining genome integrity and cell viability. This implies that targeting MTH1 activity could form the basis of a new targeted therapy strategy [1, 28]. However, other recent data challenged these observations and conclusions. In these studies, MTH1 deficiency did not hinder the growth of HeLa, SW480 or U2OS cells, and highly specific MTH1 inhibitors displayed only weak cancer cell cytotoxicity [29-31]. The title of the latest review on the topic, “MTH1 as a chemotherapeutic target: the elephant in the room” [32], highlights the fact that the disagreements remain unresolved. Hence, it has become critically important to undertake further work to shed light on these contradictory findings and better understand the relevance of MTH1 in cancer and therapy.

Lung cancer is the leading cause of cancer death worldwide [33]. Despite improvements in survival rates for many other cancer types in recent years, NSCLC therapy responses and patient outcomes have not significantly improved [33, 34]. In our study, we addressed two main objectives to enable us to assess the potential of MTH1 inhibition as a NSCLC targeted therapy strategy. First, we assessed if MTH1 deficiency alone is genotoxic or cytotoxic to several lung cell lines, and whether these effects were highly selective to NSCLC cells relative to normal cells. Second, we evaluated potential new combination therapy strategies by testing if targeting MTH1 enhanced the effects of current therapeutic agents. Thus, we tackle the currently opposing and contentious opinions on the significance of MTH1 in cancer biology and therapy [1, 35].

## Methods

### Cell lines and chemicals

A549 (CCL-185), H522 (CRL-5810), H23 (CRL-5800) and MRC-5 (CCL-171) cell lines were purchased fully authenticated from ATCC. MRC-5 and A549 cell lines were cultured in EMEM (ATCC) and DMEM-high glucose (Invitrogen) media respectively, and H522 and H23 cells in RPMI 1640 medium ATCC modification. All media were supplemented with 10% (v/v) FBS (Invitrogen). Cells were cultured at 37°C in humidified atmosphere (95% air / 5% CO_2_) and maintained at a low passage by not passaging beyond 6 months’ post resuscitation. Cisplatin, gemcitabine, etoposide, phleomycin, hydroxyurea and MTH1 small molecule inhibitors (TH287, TH588) were purchased from Sigma-Aldrich.

### siRNA transfections

Performed using DharmaFECT 1 reagent (GE Healthcare) according to manufacturer’s instructions. Briefly, a transfection complex was prepared by incubating together for 20 minutes at room temperature, 7.5 μl DharmaFECT 1 reagent, 125 μl Opti-MEM medium and siRNA diluted in 125 μl Opti-MEM medium (final siRNA concentrations were 20 or 15 nM for H522 and remaining cell lines respectively). 3 X 10^5^ cells were plated with the transfection complex and incubated for 24 hours in Opti-MEM medium, after which the transfection media was replaced with standard medium. MTH1-siRNA oligonucleotide 5´->3´ sequences (Life Technologies) were sense CAUCUGGAAUUAACUGGAUtt and antisense AUCCAGUUAAUUCCAGAUGaa. Silencer Select Negative Control 1 siRNA (ThermoFisher Scientific) was used as scramble siRNA control.

### Modified alkaline comet assay

DNA damage was assessed using Formamidopyrimidine-DNA glycosylase (Fpg)-modified comet assay that is a slight modification of the original method [36]. Briefly, slides containing cells embedded in 0.6 % low melting point agarose were incubated overnight at 4°C in lysis buffer (2.5 M NaCl, 100 mM Na_2_EDTA, 10 mM Tris-base, 1 % Triton X-100, pH 10). Lysed cells were treated with Fpg (final concentration 0.8 U/gel) for 30 minutes at 37°C, and subjected to alkaline electrophoresis in buffer containing 300 mM NaOH, 1 mM Na_2_EDTA, pH 13. Following neutralization with 0.4 M Tris-base, pH 7.5, slides were stained with 2.5 μg/ml propidium iodide (PI) and dried at 37°C. Comets were visualised at ×200 magnification using an Olympus BH-2-RFL-T2 fluorescent microscope fitted with an excitation filter of 515-535 nm and a 590 nm barrier filter, and images were captured via a high performance CCDC camera (COHU MOD 4912-5000/0000). % tail DNA was calculated using Komet software (Andor Technology). For radiation treatments, the Xstrahl RS320 X-Ray Irradiator system was used to expose agarose-embedded cells on ice (assessments of immediate DNA damage) or cells in suspension that were then cultured for 24 hours (recovery samples). TH287 and TH588 10 μM treatments were for 24 hours in complete medium.

### ROS level measurements

30,000 cells per well were seeded in triplicate in black 96 well plates (Porvair) and cultured for 24 hours. Cells were washed with 200 μl PBS prior to the addition of 1 μl H2DCF-DA (Thermo Fisher Scientific) and incubated in the dark for 30 minutes in a humidified atmosphere at 37°C. 200 μl PBS was then added to all the wells, and the relative ROS-induced fluorescence intensities were measured immediately on a FLUOstar OPTIMA Microplate Reader (BMG Labtech; 485 nm excitation and 520 nm emission wavelengths). 30-minute pre-treatment with 9.8 mM hydrogen peroxide was used for positive controls (relatively high dose used to overcome the scavenging of extracellular hydrogen peroxide by sodium pyruvate in the media [37]). Samples without seeded cells used as blanks.

### WST-1 cell proliferation assay

WST-1 is a water-soluble tetrazolium salt that is cleaved to a formazan dye in a mechanism mainly dependent on NAD(P)H production by metabolically active cells. 2 days after transfection, 1 X 10^4^ cells (2 X 10^4^ for H522) were seeded in triplicate for each sample in clear flat bottom 96 well plates, and left for 3 days before performing the assay according to manufacturer’s instructions (Sigma-Aldrich). Briefly, 10 μl Cell Proliferation Reagent WST-1 was added to each well containing 100 μl media and incubated for between 30 minutes to 4 hours. Absorbance values (that ranged between 0.5-2) were determined on an ELx808 microplate reader (BIO-TEK Instruments) at 450 nm against a blank control background. Cell proliferation (%) was determined by calculating (mean absorbance of sample / mean absorbance of control) X 100. 2-day VP-16 treatments were used as positive controls.

### Annexin V/PI apoptosis assay

Double staining with annexin V-FITC (apoptosis marker)/PI combined with flow cytometry was applied as described in manufacturer’s instructions (Affymetrix). VP-16 (etoposide) was used as a positive control, while 4 untreated negative control cells were included for instrumental compensation and gating: no stain, PI only, annexin V-FITC only, and both PI and annexin V-FITC. Samples were analysed on a BD FACSCanto™ II flow cytometer (BD Biosciences) using BD FACSDiVa™ v6.1.3 software. At least 10,000 events were acquired per sample.

### Western blot analysis

Standard techniques were used. Briefly, protein samples were prepared using Laemmli buffer lysis and sonication (15 seconds at 14 μm using Soniprep 150). MTH1, MTH2 and α-tubulin antibodies were purchased from Abcam, and CHK1 (2G1D5), phospho-CHK1 (Ser345), phospho-CHK2 (Thr68) from Cell Signaling Technology. Polyclonal secondary antibodies were horseradish-peroxidase-conjugated, and detection was performed using ECL substrate (Pierce) and X-ray film (CL-XPosure). Band intensities were quantified using densitometry GeneSnap or Image J 1.49 version software.

### Data analysis and statistical tests

GraphPad Prism software (version 7 and 6.05) was used to calculate mean ± standard deviation (S.D) or standard error mean (SEM). Unless otherwise stated in the figure legend, data was evaluated by one-way ANOVA followed by post-hoc Tukey’s multiple comparison test to compare values between two or more groups. P-value < 0.05 was considered as statistical significant. The number of independent experimental repeats are indicated in each figure or figure legend.

## Results

### MTH1 deficiency increases levels of DNA oxidation in NSCLC cells but not normal lung fibroblasts

In order to observe the effects of MTH1 deficiency we used siRNA to successfully knockdown MTH1 in various lung cell lines. We targeted MTH1 in H23 adenocarcinoma (p53-mutated), H522 adenocarcinoma (p53-mutated) and A549 lung carcinoma (wildtype p53) NSCLC cell lines in order to assess if any observed effects are applicable to NSCLC in general or are dependent on p53 status, and in MRC-5 normal lung cells to determine if the effects were cancer specific. MTH1 levels were significantly reduced 3 days after MTH1 siRNA transfection, averaging decreases of 88% in H23, 83% in A549 and 76% in MRC-5, and remained low for at least 6 days (Fig. 1A, 1B and 1D). MTH1 knockdown efficiency was slightly delayed in H522 cells, but MTH1 levels still decreased by 70% inhibition by day 4 (Fig. 1C). The levels of MTH1 depletion were comparable to those in previous published studies [11, 38]. Hence, using siRNA we were able to study the role of MTH1 in various NSCLC cell lines and normal lung cells. The level of another Nudix hydrolase family enzyme member, MTH2, did not increase following MTH1 knockdown (Additional file 1: Figure S1), suggesting it does not act to compensate for loss of MTH1. This is concordant with a recent study that found MTH2 has a preference for nucleotide substrates different to those of MTH1 [39].

**Fig. 1.**
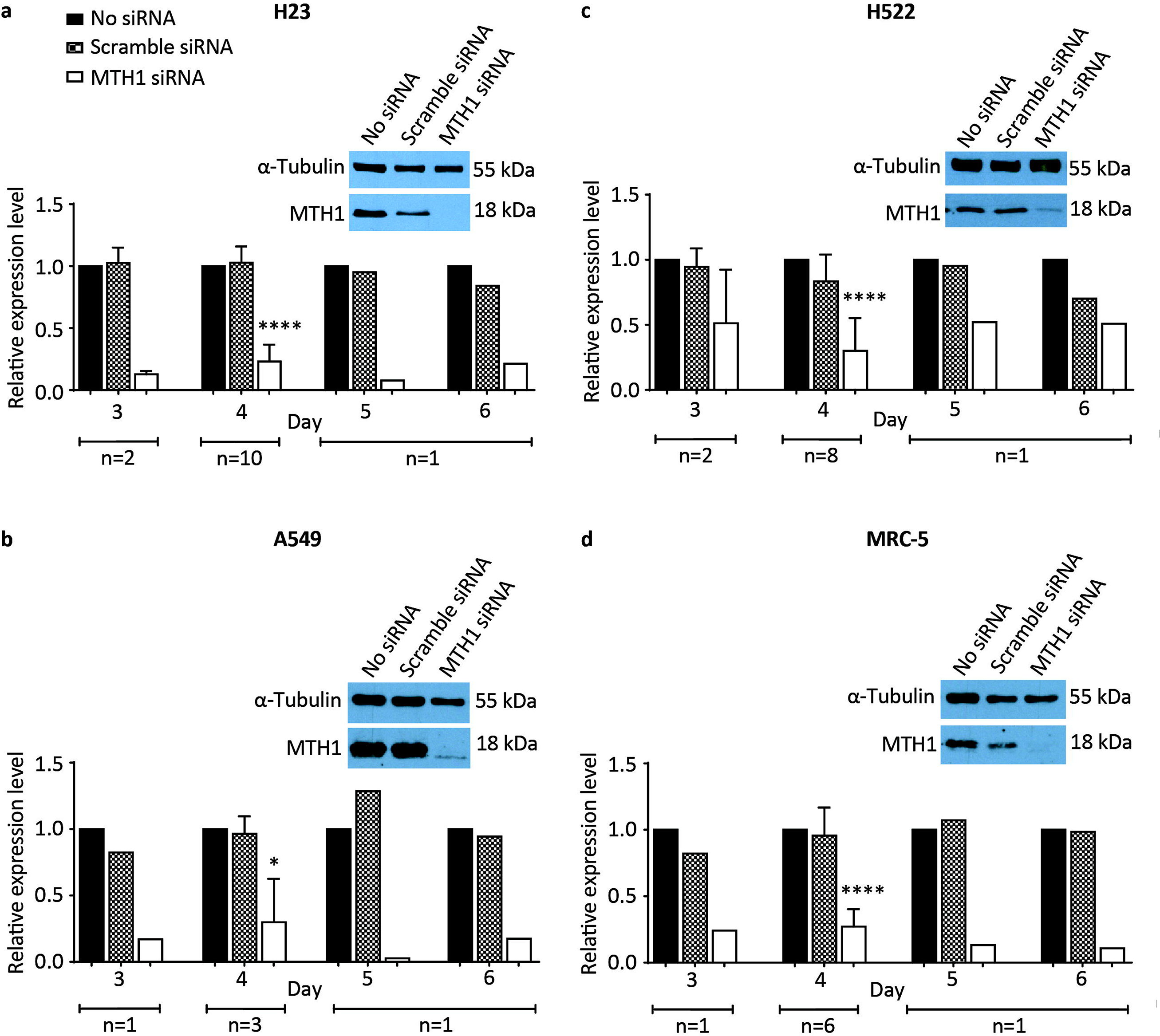
MTH1 is efficiently knocked down in various lung cancer cell lines and normal lung fibroblasts. Western blots to determine MTH1 protein levels in cell cultures grown in media without transfection reagent (no siRNA), or following transfection with MTH1 siRNA or scramble siRNA. **a** H23. **b** A549. **c** H522. **d** MRC-5. Representative day 4 blots shown. Day 4 MTH1 band intensities were normalized to corresponding α-Tubulin loading control bands, and then siRNA samples were normalised to corresponding no siRNA bands. Numbers of independent experiments (n) are indicated. Mean values and SD were calculated from the normalised values of the independent experiments. Error bars represent SD. Asterisks represent a significant difference between MTH1 siRNA and corresponding no siRNA normalised values (****P<0.0001, *P<0.05).

To begin to determine if significantly reduced MTH1 levels lead to functional consequences in different lung cell lines, we first assessed the effects of MTH1 knockdown on DNA oxidation levels using the Fpg-modified alkaline comet assay (Fig. 2A). Cells were collected 4 days after siRNA transfection to provide sufficient time for them to have undergone DNA replication in the presence of notably reduced MTH1 levels. Following MTH1 knockdown, we consistently observed significant 1.5- to 2-fold increases in oxidatively damaged DNA bases (i.e. Fpg-sensitive sites) in H23, H522 and A549 genomic DNA relative to the scrambled siRNA controls, but no difference in MRC-5 (Fig. 2B to 2E). This finding is concordant with the notion that MTH1 acts to suppress the misincorporation of oxidised dNTPs specifically in cancer cells. We observed no increases in DNA single-strand break levels (SSBs; i.e. Fpg-independent sites), which contrasts with a previous observation in A549 cells [25]. The reason for this contradiction is unclear; however, unless a cell line also harbours a base excision repair defect that leads to the generation of more or longer-lived SSB intermediates following initial damaged base removal [40], we would not predict an increase in SSB levels following MTH1 knockdown.

**Fig. 2.**
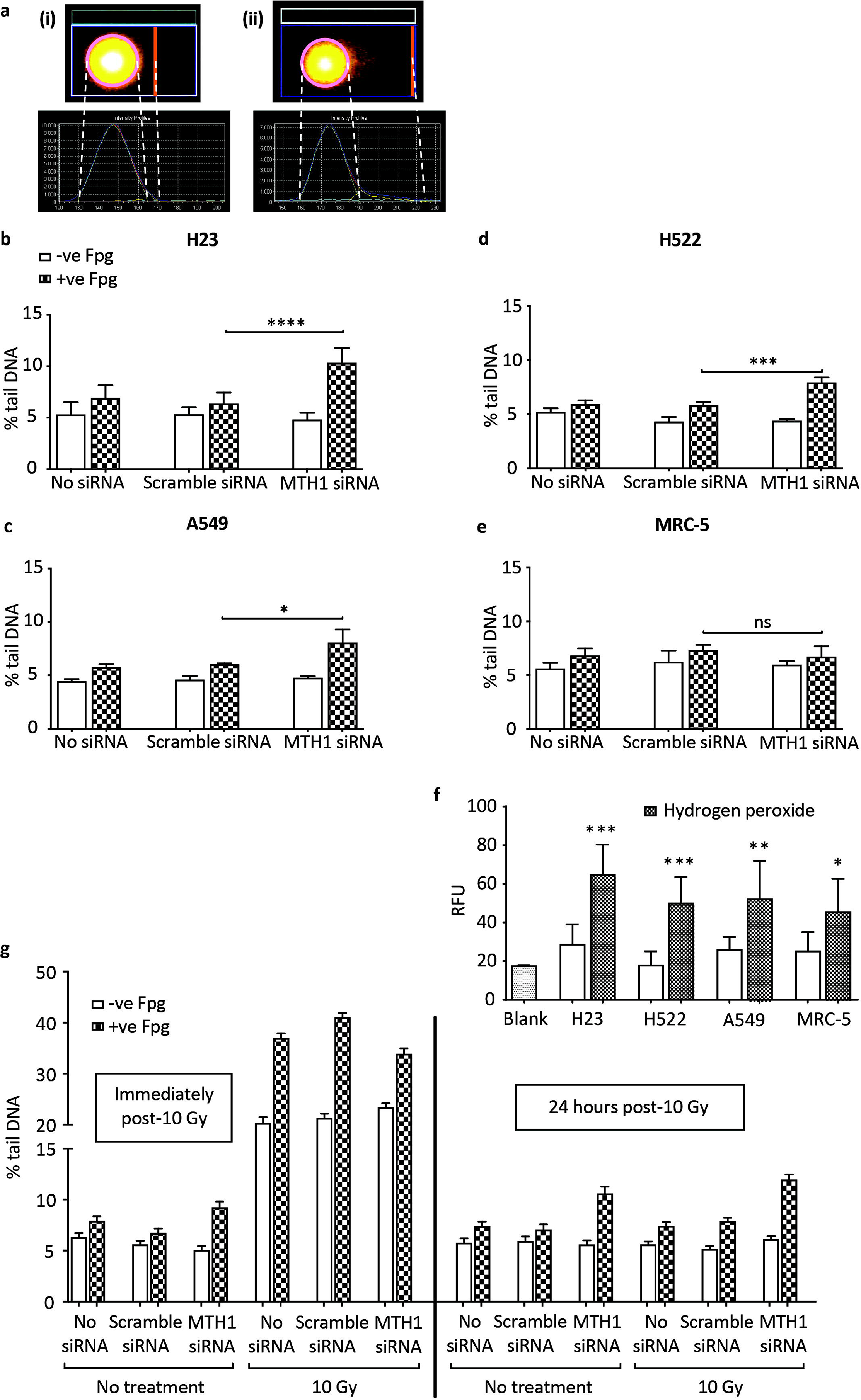
MTH1 knockdown leads to increased oxidised DNA base levels in lung cancer cell genomes. Formamidopyrimidine-DNA glycosylase (Fpg)-modified alkaline comet assay to determine DNA damage levels (expressed as % tail DNA) in individual cells grown in media without transfection reagent (no siRNA), or 4 days after transfection with MTH1 siRNA or scramble siRNA. DNA single-strand breaks detected as Fpg-independent signal, while oxidatively damaged DNA bases detected by treatment with Fpg. a Representative images of “Comets” and the corresponding intensity profiles, showing (i) ∼5% Tail DNA damage, typical of the NSCLC cells treated with no siRNA, scramble siRNA or NUDT1 siRNA and analysed by regular alkaline comet assay; and (ii) comets showing ∼10% tail DNA, typical of the NSCLC cells treated with NUDT1 siRNA and analysed by Fpg-modified alkaline comet assay (0.8U Fpg/gel). Superimposed on the Comet images are the image analysis software (Komet 5.5, Andor Technology) determined boundaries demarcating the ‘Comet head’ (pink circle) and ‘tail extent’ (vertical orange line). % tail DNA = 100 - % head DNA; % head DNA = (integrated optical head intensity / (integrated optical head intensity + integrated optical tail intensity)) x 100. b H23, 8 independent experiments. c A549, 3 independent experiments. d H522, 4 independent experiments. e MRC-5, 3 independent experiments. For (b) to (e), 200 randomly selected individual comets were scored for each sample per experiment. Mean values from independent experiments were used to generate final mean values and SD. Error bars represent SD. Asterisks indicate a significant difference between Fpg-treated MTH1 siRNA and scramble siRNA experimental means (****P<0.0001, ***P<0.001, *P<0.05); ns, not significant. f Internal ROS levels determined by measuring fluorescence signal induced by 2’,7’-dichlorodihydrofluorescein diacetate oxidisation. RFU, relative fluorescence units. Blank samples were without seeded cells. Mean values were calculated from 4 independent experiments. Error bars represent SD calculated from the independent experiment values. Unpaired T-test was performed. Asterisks indicate a significant difference between hydrogen peroxide treated and untreated samples (***P<0.001, **P<0.01 and *P<0.05). g Comet assay post-irradiation of H23 cells. Error bars represent SEM calculated from 400 individual comet values analysed in total from 2 independent experiments.

To assess if the conditional requirement for MTH1 is due to cancer-associated oxidative stress we measured ROS levels in all cell lines. However, we found no differences between any of the NSCLC lines and MRC-5 (Fig. 2F). This is consistent with there being no significant differences between any of the cell lines in the background levels of DNA oxidation and SSBs (Fig. 2B to 2E), but implies that the basis of the NSCLC requirement for MTH1 to suppress DNA oxidation in NSCLC cells is not simply due to higher ROS levels. This finding goes against current proposed models [22, 25]. Given all the genetic and signaling disturbances within cancer cells, there may be many other causes of this MTH1 dependency that remain to be discovered.

### MTH1 is not required in response to exogenous sources of oxidative stress

Hydrogen peroxide treatment was previously shown to lead to the accumulation of genomic 8- oxoGua and cell death in *MTH1*^-/-^ mouse embryonic fibroblasts [41], indicating that oxidative stress can be cytotoxic in a MTH1-deficient background. We proposed that in addition to a role in processing oxidised dNTPs that are generated endogenously within NSCLC cells, MTH1 would also be required to suppress the misincorporation of damaged DNA bases following exposure to exogenous sources of oxidative stress and cancer therapy agents. To determine this, we first assessed whether higher DNA oxidation levels were detectable in MTH1-deficient H23 cells after irradiation (IR) treatment, which targets the nucleotide pool [42]. Cell samples were analysed immediately after IR and following a 24-hour recovery, which was permitted to allow enough time for IR-generated oxidised dNTPs to be misincorporated. The relative increases in SSB levels and oxidatively damaged DNA immediately after IR did not differ between the scramble siRNA control and MTH1-deficient cultures (Fig. 2G), confirming that MTH1 does not have a role in preventing direct oxidation of DNA. However, by 24 hours post-IR, the relative levels of oxidatively damaged DNA in all samples had returned to levels comparable to those prior to IR. A similar observation was seen when oxidative stress was induced after treatment with the model oxidant (non-radical ROS), hydrogen peroxide (Additional file 2: Figure S2). Overall, this suggests that MTH1 is not required to prevent the misincorporation of dNTPs that are oxidised via exogenous agents. Alternatively, other MTH1- independent compensatory factors such as Ogg1 may be activated when very high levels of damaged dNTPS are acutely generated [43].

### MTH1 deficiency induces alterations in DNA damage response signaling

We propositioned that the increased levels of oxidised DNA bases caused by MTH1 knockdown may lead to DNA replication stress in NSCLC cell lines, while normal cells would remain genomically stable. The central kinase pathways in the DNA-replication-associated DDR are ATR-CHK1 and ATM-CHK2, which are initially activated by defective DNA replication forks and DSBs respectively [44]. Using Western blotting, we detected indications of DDR alterations in all NSCLC cells lines following MTH1 knockdown (Fig. 3), suggesting that the cells were responding to replication stress and some kind of secondary DNA damage. Surprisingly, however, the DDR responses in different NSCLC cell lines varied in the pathways affected and whether they were activated or repressed.

**Fig. 3.**
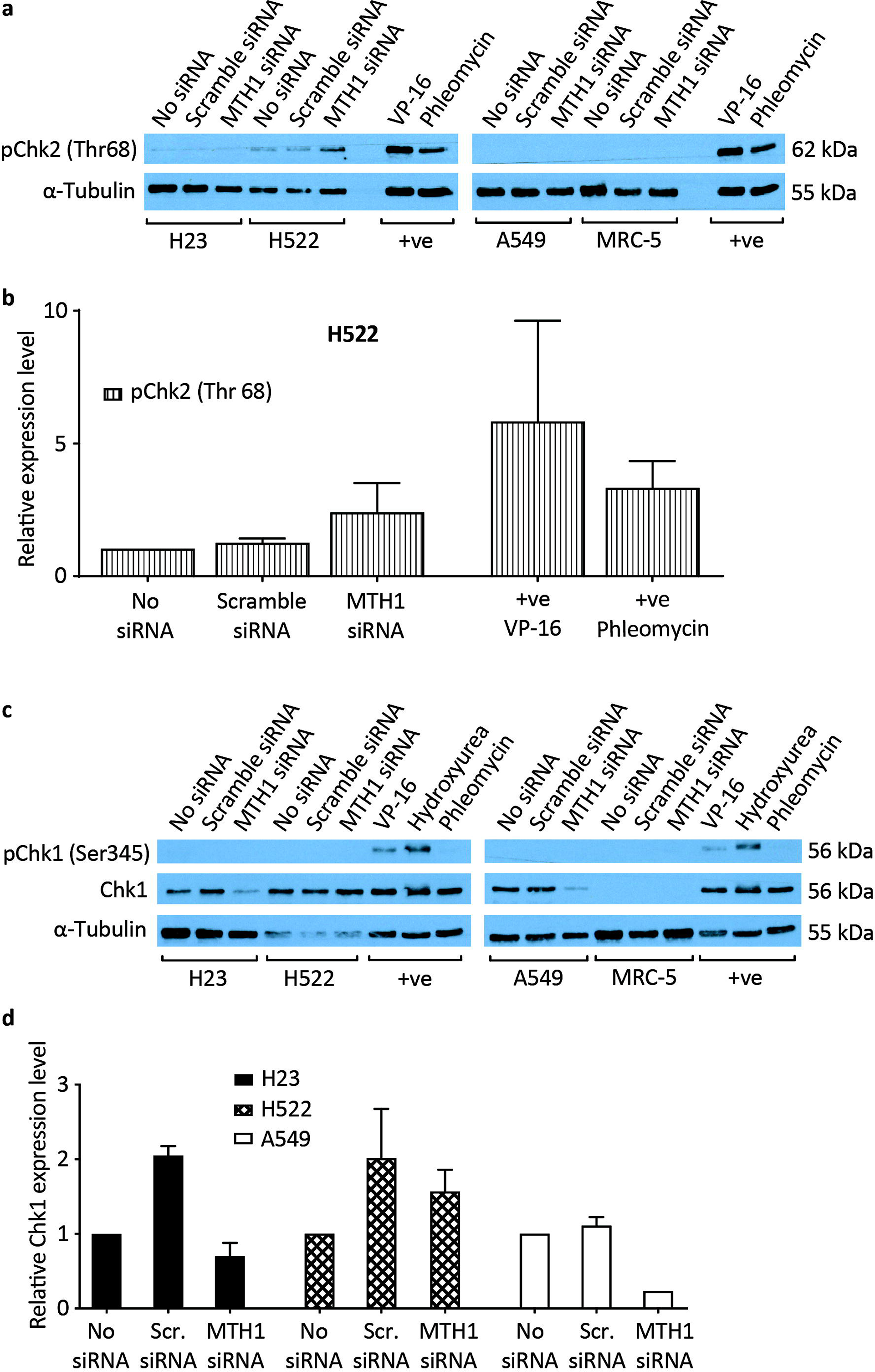
Alterations in DNA damage response signalling following MTH1 knockdown. Cells were grown in media without transfection reagent (no siRNA), or transfected with MTH1 siRNA or scramble siRNA (Scr. siRNA). Western blots were performed 4 days post-transfection. Positive control samples (+ve) were H23 cells treated with VP-16 (etoposide, 25 μg/ml), phleomycin (25 μg/ml) or hydroxyurea (2 mM) for 2 hours. **a** and **c** Representative Western blots. **b** pChk2(Thr68) band intensities from H522 samples were normalised to α-Tubulin, and expression levels calculated relative to no siRNA samples. **d** Chk1 Western blot band intensities were normalized to α-Tubulin, and expression levels calculated relative to no siRNA samples. Mean values and SD were calculated from the normalised values of the 3 independent experiments. Error bars represent SD. Asterisks represent a significant difference between MTH1 siRNA and no siRNA normalised values (****P<0.0001).

We detected DDR activation in MTH1-knockdown H522 cells, as indicated by an approximately 2-fold increase in CHK2 phosphorylation levels relative to no siRNA and scramble siRNA controls (Fig. 3A and 3B). This is indicative of the presence of DSBs, as shown by use of the topoisomerase ll inhibitor VP-16 (etoposide) as a positive control. In contrast, we repeatedly detected notable losses of CHK1 protein levels in MTH1-knockdown H23 and A549 cells relative to no siRNA and scramble siRNA controls, which was significant in A549 relative to no siRNA, but no indications of increased CHK1 phosphorylation in any cell line (Figures 3C and 3D). The basis of the CHK1 down-regulation is not clear. It may indicate that MTH1-deficient H23 and A549 cells had increased DNA replication stress levels, but that there was a selective pressure to rapidly adapt and down-regulate associated ATR- CHK1 activity to overcome growth arrest. Transfection with the scrambled siRNA caused increased Chk1 levels in H23 and H522 cells. The reason for this is unclear, but it is very likely due to unavoidable transfection-dependent siRNA-independent effects on these particular cell lines as the commercially available scrambled siRNA used (Silencer Select Negative Control No. 1, ThermoFisher Scientific) has no significant sequence similarity to human gene sequences and is confirmed to have minimal effects on gene expression. Nonetheless, the fact that lower Chk1 levels were detected in H23 and A549 MTH1 siRNA cultures even relative to the basal no siRNA (no transfection) cultures demonstrates the strength of the phenotype. Finally, in our experimental conditions we could not detect total CHK1 in MRC-5 cells (ATCC CCL-171), consistent with a previous study [45]. This may be due to very low relative expression levels and/or an issue with the particular antibody used.

### MTH1 promotes H23 cell proliferation, but is dispensable for NSCLC cell survival

Given the observed function for MTH1 in genome maintenance, we hypothesised that MTH1 would be ‘conditionally essential’ in NSCLC. To first test this, we assessed if MTH1 was required for NSCLC cell proliferation using the WST-1 assay, which measures the metabolic activity in cell cultures. Concordant with the indication in Fig. 3 that H23 and H522 cell lines are slightly sensitive to the non-specific effects of transfection, the scrambled siRNA transfected cultures showed 20 % and 30 % decreases in cell proliferation, respectively (Fig. 4A and 4C). Nevertheless, MTH1 knockdown induced a significant 54 % and 34 % decrease in H23 cell proliferation relative to the no siRNA and scrambled siRNA controls, respectively (Fig. 4A), indicating that MTH1 is partially required for H23 growth. However, similar decreases in cell proliferation were not seen in A549, H522 or MRC-5 cells relative to controls (Fig. 4B to 4C). This contradicts previous data that suggested MTH1-deficient A549 cells have dramatic proliferation defects [25].

**Fig. 4.**
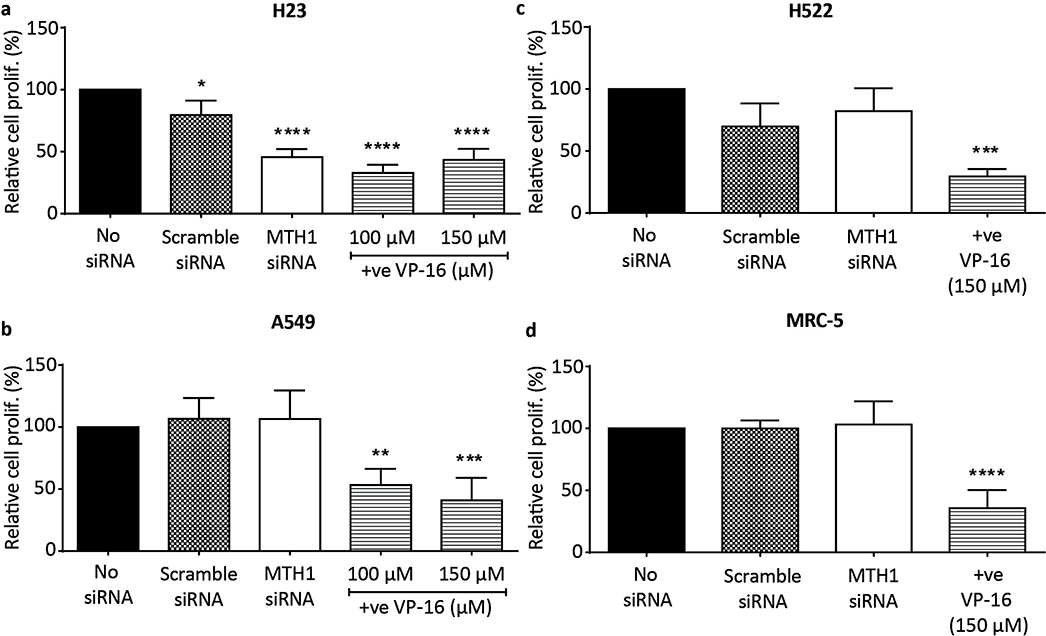
MTH1 targeting reduces H23 cell proliferative capacity. WST-1 assay on cells grown in media without transfection reagent (no siRNA), or 5 days after transfection with MTH1 siRNA or scramble siRNA. **a** H23. **b** A459. **c** H522. **d** MRC-5. (a), (b) and (d) Values from 4 independent experiments were used to generate final mean values and SD. (c) Values from 3 independent experiments were used to generate final mean values and SD. Error bars represent SD. Asterisks indicate a significant difference relative to corresponding no siRNA controls (****P<0.0001, ***P<0.001, **P<0.01, *P<0.05).

To determine if MTH1 deficiency in NSCLC induces cell death in addition to or rather than cell growth inhibition, we measured apoptosis levels using annexin V (Fig. 5A). We did not observe increased apoptosis levels in any MTH1 knockdown cell cultures relative to the scrambled siRNA controls irrespective of the p53-status of the line (Fig. 5B to 5E). Consistent with previous observations (Fig. 3 and 4), H23 and H522 scrambled siRNA cultures demonstrated minor MTH1-independent transfection-dependent effects. The results of the apoptosis assay were confirmed using another MTH1 siRNA that induces a similar decrease in MTH1 levels (Additional file 3: Figure S3).

**Fig. 5.**
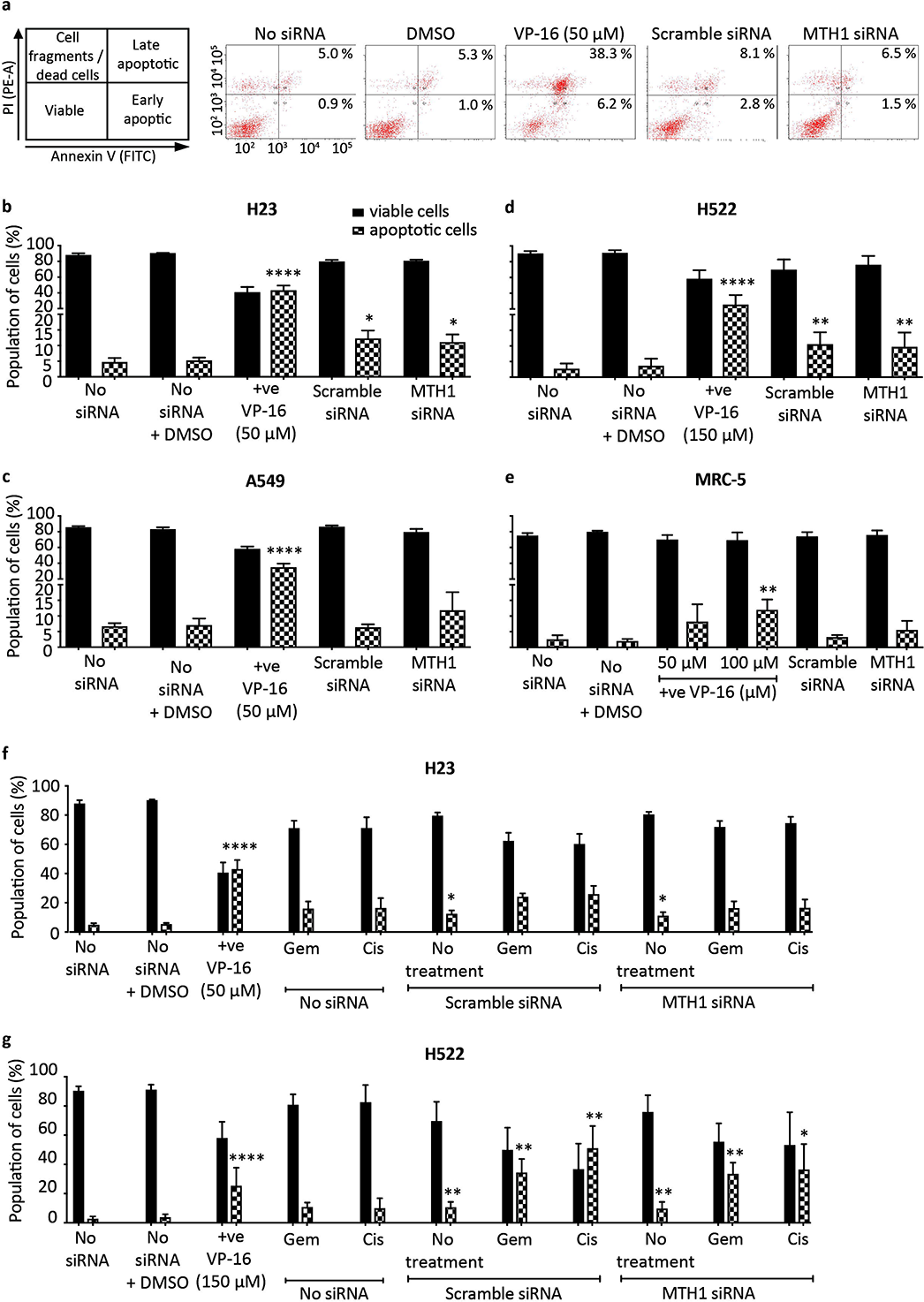
MTH1 deficiency does not induce apoptosis or augment the cytotoxic effects of chemotherapy agents. Apoptosis assay to determine viability of cells cultured for 4 days in media without transfection reagent (no siRNA), or following transfection with MTH1 siRNA or scramble siRNA. Positive control of 48 hours VP-16 treatment (+ve) also included. Harvested cells were dual stained with annexin V-FITC/propidium iodide (PI) and assessed by flow cytometry. Annexin V is an apoptosis marker. PI is a DNA stain that is excluded from viable and early apoptotic cells. Percentage values from independent experiments were used to calculate final mean values and SD. Error bars represent SD. **a** Representative bivariate plots of H23 cells. **b** H23. **c** A549. **d** H522. **e** MRC-5. **(b)** and **(c)** 3 independent experiments performed. **(d)** and **(e)** 4 independent experiments performed. Asterisks indicate a significant difference relative to corresponding no siRNA control (or no siRNA + DMSO control in case of VP-16). **f** H23 cells. 2 days after transfection, 0.01 μM gemcitabine (Gem) or 5 μM cisplatin (Cis) were added to the appropriate cultures for the remaining 48 hours (0.5% DMSO). 3 independent experiments performed with siRNA transfections (5 repeats for non-transfected samples). **g** H522 cell line. 2 days after transfection, 40 μM gemcitabine (Gem) or 10 μM cisplatin (Cis) were added to the appropriate cultures for the remaining 48 hours (1.5% DMSO). 3 independent experiments performed with siRNA transfections and Cis and Gem treatments (4 repeats for untreated and non-transfected samples). **(f)** and **(g)** Asterisks in Scramble siRNA and MTH1 siRNA samples indicate a significant difference between No treatment, Gem or Cis and corresponding no siRNA + DMSO, no siRNA + Gem and no siRNA + Cis percentage values, respectively (****P<0.0001, **P<0.01, *P<0.05).

Though MTH1 inhibition alone did not cause apoptosis, we propositioned that it would still enhance the targetting and effectiveness of current chemotherapy agents that induce oxidative stress [46, 47] and DNA replication stress. The basis of this idea was that MTH1 inhibition leads to higher levels of oxidatively damaged DNA in NSCLC cells but not normal cells, which when combined with therapy-induced effects selectively pushes the DNA damage levels over the cytotoxic threshold in NSCLC cells. In particular, the mechanisms of action of gemcitabine and cisplatin suggests the combining them with the effects of MTH1 inhibition could lead to additive or synergistic effects that would improve patient outcomes. Gemcitabine is a chemical antimetabolite and analogue of deoxycytidine that induces DNA replication stress [48-50], and cisplatin leads to DNA replication defects via the formation of DNA crosslinks [51, 52] and can increase intracellular ROS levels possibly through mitochondrial dysfunction and activation of NAD(P)H oxidase and superoxide production [53-55].

We treated H23 and H522 MTH1 knockdown cells with gemcitabine or cisplatin and monitored apoptosis levels. The effect on H522 was of particular interest, as relative to H23 and A549, this cell line displays much higher resistance to etoposide, gemcitabine and hydrogen peroxide (Fig. 5 and Additional file 4: Figure S4). Despite our predictions, a combination of MTH1 knockdown with either gemcitabine or cisplatin treatment did not lead to significant increases in cell death levels relative to the treated scramble siRNA controls (Fig. 5F and 5G). These results were reproducible with a different MTH1 siRNA (Additional file 3: Figure S3). This demonstrates that the oxidative DNA damage induced by MTH1 deficiency in NSCLC cells does not sufficiently sensitise them to the effects of current therapeutic agents.

### TH287 and TH588 MTH1 inhibitors have variable effects on NSCLC cell lines

Using siRNA to assess the effects of MTH1 deficiency on lung cell lines had uncovered some observations that agreed with our predictions, but also uncovered some unexpected results. To confirm our findings, we similarly used the small molecule MTH1 inhibitors, TH287 and TH588, which were previously shown to lead to a dramatic increase in oxidatively damaged DNA and loss of viability selectively in cancer cells [22, 38].

When NSCLC and MRC-5 cell lines were treated with the same dose (10 μM) of TH287 and TH588, an observable though not significant increase in the levels of oxidatively damaged DNA was evident only in H23 cells (Fig. 6A to 6D), and there was no increase in SSB levels in any cell line. This implies that TH287 and TH588 treatments do not lead to increased in oxidised dNTP misincorporation in H522, A549 or MRC-5 cells. We next assessed if the levels of oxidatively damaged DNA correlated apoptosis induction. Indeed, TH287 and TH588 induced significant 3.2 and 3.0-fold increases in apoptosis levels in H23 cells, respectively (Fig. 6E), and did not affect A549 cells (Fig. 6F). However, significant 2.8 and 3.5-fold increases in apoptosis levels were also observed in inhibitor-treated H522 cells (Fig. 6G), indicating that levels of DNA oxidation were not the major basis of the cytotoxic effects. Normal MRC-5 lung fibroblasts did not exhibit any apoptotic response to TH287 and TH588 (Fig. 6H), agreeing with the original suggestion that TH287 and TH588 are not cytotoxic to normal cells [22]. Collectively, these data suggest that TH287 and TH588 at the dose used induce cell death in p53-deficient cancer cells through ‘off target effects’ rather than inhibition of MTH1, but do not induce cell death in p53-proficient NSCLC cells and normal cells.

**Fig. 6.**
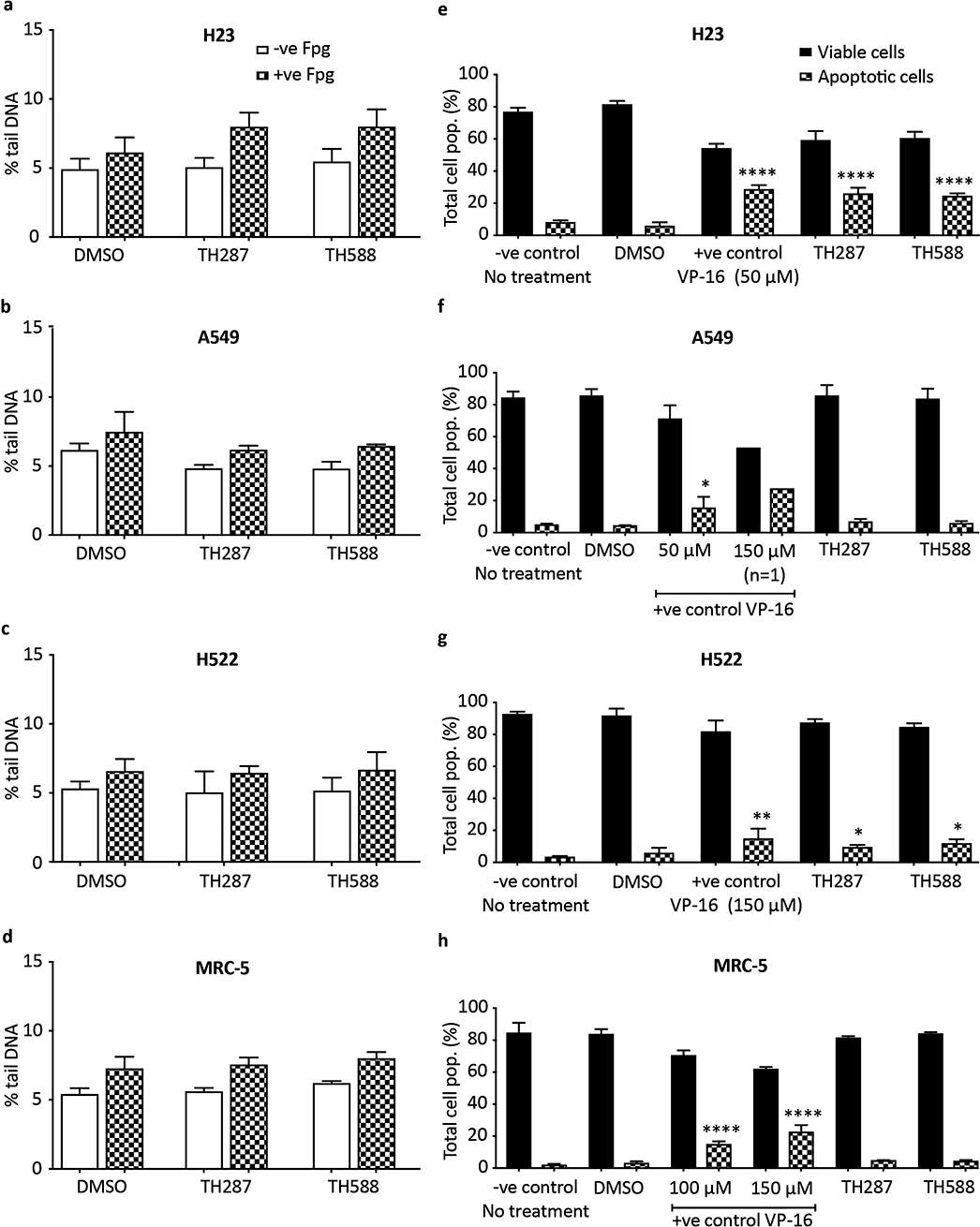
Variable effects of TH287 and TH588 MTH1 inhibitors on DNA oxidation and apoptosis. **a** to **d** Fpg-modified alkaline comet assay. DMSO (0.066 % v/v) used as a vehicle control. Means ± SD calculated from 3 independent experiments. 200 randomly selected individual comets were scored for each sample per experiment. Mean values calculated from 3 independent experiments were used to generate final mean values and SD. Error bars represent SD. **e** to **h** Annexin V-FITC/PI apoptosis assay to determine cell viability. DMSO (0.5-1.5% v/v) or VP-16 (+ve) were applied as vehicle controls and positive controls, respectively. 3 independent experiments performed (except 150 μM VP-16). Percentage values from each experiment were used to calculate final mean values and SD. Error bars represent SD. Asterisks indicate a significant difference between treated and untreated percentage values.

## Discussion

In this study we tested the potential of a new targeted therapy strategy for NSCLC, whilst simultaneously analysing opposing opinions within the field regarding the conditionally essential requirements for MTH1 in cancer cells and whether the current pursuit of MTH1 inhibitor development is likely to yield effective therapeutic agents [1, 32, 35]. We show that MTH1 does indeed have a NSCLC-specific role for maintaining genome stability. The basis of this cancer-specificity remains unclear, as DNA oxidation levels in MTH1-deficient lung cells do not correlate with background ROS levels. This goes against the current model [32], and suggests that perhaps the NSCLC-specific effect could be due to downstream defects in removing the oxidatively damaged DNA induced in the cancer cell lines [56, 57]. Despite the functional role for MTH1 in NSCLC cells, we show that MTH1 deficiency ultimately does not cause NSCLC death, either alone or when combined with other therapeutic agents. One possibility for this could have been that the cell culture media used contained sodium pyruvate, a ROS scavenger. However, we do not believe this to be the case as sodium pyruvate scavenges extracellular ROS rather than intracellular endogenous ROS [37, 58], and from what we can tell, other MTH1 studies that reported cytotoxic effects associated with MTH1 also used media containing sodium pyruvate [11, 22, 29]. Ultimately, our work argues that MTH1 inhibitors will likely not be effective therapeutic agents. Instead, given that we show that MTH1 deficiency in NSCLC cells induces non-cytotoxic DNA oxidation and DDR alterations, we propose that treating NSCLC patients with MTH1 inhibitors could actually provide an environment for further mutation accumulation to drive cancer heterogeneity and evolution. In accordance with this proposition, MTH1-knockdown in human B lymphoblastoid cells induces a higher mutation rate but not cell death after UVA-induced oxidative stress [59], while MTH1 overexpression repressed the DNA-replication-dependent mutator phenotype in mismatch-repair-defective colorectal cancer cells [60].

The increases in oxidatively damaged DNA in NSCLC cell lines following MTH1 knockdown was relatively small (Fig. 2). However, the alterations in DDR signaling indicate that this was enough to disrupt DNA replication and lead to secondary types of DNA damage such as DSBs (Fig. 3). One proposed model for how this occurs is that oxidised DNA bases induce DNA replication stress, which is defined as defective DNA replication fork progression [61, 62], and that this somehow subsequently leads to DSBs [22, 27]. It is possible that DSBs can arise from replication fork run-off at BER-induced SSBs, which would lead to the generation of one-ended broken DNA replication forks in a mechanism analogous to DSBs arising from Top1-DNA adducts [63]. Alternatively, DNA replication forks may stall at sites of oxidatively damaged DNA and be cleaved by endonucleases to also generate one-ended broken replication forks [64, 65]. No matter how they arise, DNA DSBs are potentially highly genotoxic or cytotoxic due to loss of chromosome integrity. In particular, one-ended DSBs on replicating chromosomes may be very difficult to resolve and are linked to various types of mutations [66-69].

It is unclear why the DDR signaling alterations varied between the MTH1-defective NSCLC lines, but given that different cancers already harbour many other mutations and potentially DDR defects, the signaling variances may simply reflect the differing abilities and deficiencies in DDR functions in different cancers. Furthermore, ultimately the ATM/CHK2 and ATR/CHK1 pathways are interlinked, as ATM-activating DSBs can subsequently lead to ATR activation if they are resected to generate ssDNA overhangs [70], and processing of ATR-activating stalled forks can generate DSBs [64]. The induction of phosphorylated CHK2 in H522 cells suggests that DNA DSBs arise, but the reasons for decreased total CHK1 levels in A549 and H23 cell cultures were surprising. Although DDR activation following MTH1 knockdown was previously observed, including phosphorylation of H2AX, 53BP1, ATM and DNA-PKcs [11, 22, 24, 27], to our knowledge this is the first time that a MTH1-knockdown-associated ‘switching off’ of the DDR has been discovered. Induced deficiency of a DDR factor may indicate that MTH1 knockdown in A549 and H23 cells initially induces DNA replication fork stalling and ATR/CHK1 activation, but that the bulk of the cells efficiently turned off this cell cycle checkpoint signaling by lowering CHK1 levels to continue proliferating. Given that CHK1 levels were decreased within 4 days of MTH1 knockdown, this would not be enough for a mutation and clonal expansion to occur within the population, suggesting CHK1 suppression occurs through another mechanism that may involve changes in gene expression (epigenetic), RNA processing, post-translational modifications and/or proteosomal degradation. Accordingly, various stresses have previously been linked to CHK1 degradation [71, 72].

There have been several contradictory and opposing reports on the cytotoxicity of MTH1 deficiency using various siRNA and shRNA sequences, cell lines and inhibitors [11, 22, 24, 26, 27, 29-31, 38], which were recently summarized and compared in a review article [32]. A critical finding of our work is, that despite MTH1 deficiency causing genomic instability in NSCLC cells and decreased H23 cell proliferation, there was a lack of cytotoxicity associated with MTH1 knockdown in all NSCLC cell lines (Fig. 5). A simple explanation for this finding is that the levels of MTH1 knockdown in our experiments were not sufficient enough to induce MTH1 deficiency. Or, as already discussed, other factors may be able to sufficiently compensate for MTH1 [43]. However, we do not believe either of these possibilities is the basis of the disparities, as the 1.5- to 2-fold increase in oxidatively damaged DNA damage (Fig. 2) is comparable to that in other studies that did detect loss of cancer cell viability [11, 22, 26, 38]. Also, we performed our experiments 4 days after transfection, which was before MTH1-proficient cells could take over culture (as confirmed by Western blot, Fig. 1). Hence, we suggest that the increased levels of genomic instability in MTH1-deficient NSCLC cells is not sufficiently high enough to induce cell death, rather it could promote further mutations and heterogeneity. Overall, our data therefore indicates that MTH1 inhibition will likely not be a successful therapeutic strategy for many NSCLC patients even when used in combination treatments. However, it remains possible that the effects of MTH1 deficiency vary considerably depending on circumstances. For example, MTH1 inhibition may be more effective on cancer cells that exhibit very high oxidative stress or particular mutations, and combining MTH1 inhibition with other specific agents or inhibitors (for example, Chk2 inhibitors) may prove to be selectively toxic.

The MTH1 small molecule inhibitors, TH287 or TH588, were proposed to be effective for cancer cell killing due to MTH1 inhibition [22, 27]. In our studies, TH287 and TH588 did induce apoptosis in 2 out of the 3 NSCLC cell lines tested, but this did not entirely correlate with increases in oxidatively damaged DNA levels (Fig. 6). This suggests that the effects on cell viability may have been distinct from MTH1 inhibition. Accordingly, the cytotoxicity of TH588 to melanoma cells was recently suggested to correlate to endogenous ROS levels but be independent of MTH1 [73]. It was also recently proposed that TH287 or TH588 at the dose we used exert much of their cytotoxic effects through tubulin polymerisation defects [30], though this conclusion was subsequently challenged [38]. Repeating the treatments at lower doses that do not appear to induce tubulin polymerisation such as 2 μM [30] over a longer time period may more specifically assess the consequences of MTH1 inhibition on NSCLC cells. Nonetheless, other highly specific MTH1 inhibitors were found to not be cytotoxic to cancer cells [29, 31]. These data not only support our MTH1 knockdown findings, but also strengthen the argument that MTH1 inhibitors may not make effective therapeutic agents.

## Conclusion

The importance of the MTH1 enzyme in cancer is a highly controversial topic within current cancer research and the focus of intense study. We show that MTH1 is indeed selectively required in various NSCLC cell lines to maintain genome integrity and support H23 cell proliferation. However, unexpectedly, MTH1 is ultimately not essential for NSCLC cell viability and does not alter responses to current therapeutic agents. Thus, our work indicates that MTH1 is likely not an effective therapeutic target for NSCLC. On the contrary, inhibiting MTH1 may promote further mutation accumulation and disease progression.

## List of abbreviations

DSB: DNA double-strand breaks
DDR: DNA damage response
dNTP: deoxyribonucleoside triphosphate
Fpg: Formamidopyrimidine-DNA glycosylase
PARP: poly(ADP-ribose) polymerase
MTH1: Mut T Homologue 1
NSCLC: non-small cell lung cancer
PI: propidium iodide
ROS: reactive oxygen species
8-oxoGua: 8-oxo-7,8-dihydroguanine

## Ethics approval and consent to participate

Not applicable

## Consent for publication

Not applicable

## Availability of data and materials

All data generated or analysed during this study are included in this published article [and its supplementary information files]

## Competing interests

The authors declare that they have no competing interests

## Funding

This work was funded by the Iraqi Cultural Attache (UK).

## Author contributions

SSF, GDDJ and MDE conceived and designed the experiments. HHKA performed the experiments. KMHA assisted in performing the WST-1 assay. HHKA, SSF, GDDJ and MDE analyzed the data. SSF, GDDJ, MDE and HHKA wrote the paper. All authors read and approved the final manuscript.

## Acknowledgements

We thank the Leicester Imaging Technologies facility (LITE) within the Centre for Core Biotechnology Services of the University of Leicester for use of the flow cytometers. We also thank Kamla Alsalmani for providing training to HHKA.

## Additional files

### Additional file 1: Figure S1

MTH2 levels are stable following MTH1 siRNA knockdown. Western blots from the single confirmation experiment performed to determine MTH2 protein levels in cell cultures grown in media without transfection reagent (no siRNA), or following transfection with MTH1 siRNA or scramble siRNA (n = 1).

### Additional file 2: Figure S2

MTH1 deficiency does not alter levels of oxidatively-modified DNA following hydrogen peroxide treatment. Fpg-modified alkaline comet assay to determine DNA damage levels in individual cells grown in media without transfection reagent (no siRNA), or 4 days after transfection with MTH1 siRNA or scramble siRNA. After 30 minutes hydrogen peroxide treatment at 37°C, samples were collected either immediately or allowed to recover in fresh media. Means were calculated from 100 individual comets from a single experiment. Error bars represent SEM of comet values. a H23. b H522.

### Additional file 3: Figure S3

MTH1 knockdown with another siRNA similarly does not induce apoptosis in H23 cells. a Western blots to determine MTH1 protein levels in H23 cell cultures grown in media without transfection reagent (no siRNA), or following transfection with MTH1 siRNA (ThermoFisher Scientific, S194633, oligonucleotide 5´->3´ sequences were sense UUAACUGGAUGGAAGGGAAtt and antisense AUCCAGUUAAUUCCAGAUGaa) or scramble siRNA. Representative day 4 blot shown. Day 4 MTH1 band intensities were normalized to corresponding α- Tubulin loading control bands, and then siRNA samples were normalised to corresponding no siRNA bands. Numbers of independent experiments (n) are indicated. Mean values were calculated from the normalised values of the independent experiments. Error bars represent SD. Asterisks represent a significant difference between MTH1 siRNA and corresponding no siRNA normalised values (**P<0.01, *P<0.05). b Apoptosis assay to determine cell viability of H23 cells cultured for 4 days in media without transfection reagent (no siRNA), or following transfection with MTH1 siRNA (S194633) or scramble siRNA. Harvested cells were dual stained with annexin V-FITC/PI and assessed by flow cytometry to detect both early and late apoptosis. 2 days after transfection, 0.01 μM gemcitabine (Gem) or 5 μM cisplatin (Cis) were added to the appropriate cultures for the remaining 48 hours. Positive control of 48 hours VP-16 treatment (+ve) also included (n = 1).

### Additional file 4: Figure S4

Apoptotic dose responses of H23 and H522 cell lines to various genotoxic agents. Apoptosis assay to determine cell viability following 48 hours treatment with different agents. In addition, cells were exposed to VP-16 as positive controls, and DMSO (0.5-2% v/v) and untreated negative control samples were also included. Harvested cells were dual stained with annexin V-FITC/PI and assessed by flow cytometry to detect both early and late apoptosis. a Hydrogen peroxide treatment of H23 cells. b Hydrogen peroxide treatment of H522 cells. c Gemcitabine treatment of H23 cells. d Gemcitabine treatment of H522 cells. e Cisplatin treatment of H23 cells. f Cisplatin treatment of H522 cells. Percentage values from independent experiments were used to calculate final mean values and SD. Error bars represent SD. All experiments were repeated more than 3 times. Asterisks indicate a significant difference relative to corresponding DMSO controls (****P<0.0001, ***P<0.001, **P<0.01, and *P<0.05).

